# Badnaviruses of sweetpotato: symptomless co-inhabitants on a global scale

**DOI:** 10.1101/140517

**Authors:** Jan F. Kreuze, Ana Perez, Marco Galvez, Wilmer J. Cuellar

## Abstract

Sweetpotato is among the most important root-crops worldwide, particularly in developing countries, and its production is affected severely by a variety of virus diseases. During the last decade a number of new viruses have been discovered in sweetpotatoes from different continents through next generation sequencing studies, among them belonging to the genus *Badnavirus* and collectively assigned to the species *Sweet potato pakkakuy virus* (SPPV). We determined the complete genome sequence of two SPPV isolates and show the ubiquitous presence of similar viruses in germplasm and field material from around the globe. We show SPPV is not integrated into the sweetpotato genome, occurs only at extremely low titers but is nevertheless efficiently transmitted through seeds and cuttings. They are unaffected by virus elimination therapy and lack any discernible symptoms in sweetpotatoes or indicator host plants. Nevertheless, they show considerable variation in their nucleotide sequences and correspond to several genetic lineages. Studies of their interaction with the two most important sweetpotato viruses showed only limited synergistic increase in the titres of one of two SPPV isolates. We contend that these viruses may pose little threat to sweetpotato production and more likely represent a new type of persistent virus in a possibly commensal or mutualistic relationship with sweetpotato.

**Importance:** Next generation sequencing approaches have in the last few years led to the discovery of many virus like sequences in different crop plants including sweetpotatoes. The significance of such discoveries can sometimes be elusive when they have not been associated with specific symptoms due to mixed infections or have been found in apparently healthy plants. Badnavirus sequences found in sweetpotatoes provide a typical case. Considering they have now been reported globally, it was important to determine how common these viruses are and what their possible impact may be on sweetpotato production. The significance of our research lies in resolving the case of badnaviruses, providing evidence they represent a new type of vertically transmitted persistent and apparently harmless episomal viruses living in a state of commensalism with their host.

## Introduction

Sweetpotato is one of the most important foodcrops worldwide, particularily in developing countries, where it serves as a food security crop, animal feed as well as for processing. Currently orange fleshed varieties are being promoted in sub-Saharan Africa to combat vitamin A deficiency due to their high content of pro-vitamin A. Being clonally propagated, sweetpotatoes suffer from the accumulation of viral diseases over generations, leading to reduced yields. More than 30 viruses have been reported from sweetpotato to date, with most of them belonging to the families *Potyviridae, Geminiviridae* and *Caulimoviridae* (1). The most important among the sweetpotato viruses is probably sweet potato chlorotic stunt virus (SPCSV; genus *Crininvirus*, family *Closteroviridae*), as it is able to compromise resistance of sweetpotato to other viruses causing synergistic viral diseases co-infection (2–8). The most important synergistic disease is caused by co-infection of SPCSV and sweet potato feathery mottle virus (SPFMV; genus *Potyvirus*, family *Potyviridae*) and may be exacerbated by infection with additional viruses (7, 8).

Some of the more recently discovered viruses in sweetpotato are sweet potato badnavirus A and B ((9), which have collectively been assigned to the species *Sweet potato pakkakuy virus* (SPPV, family *Caulimoviridae*, genus *Badnavirus*). Although SPPV have already been identified on all continents using various methods (10–14), little is still known about the biology of this group of viruses. Badnaviruses (15) infect a broad range of important crops including monocots and dicots, although most species have a limited host-range. They often infect perennial corps and symptoms are mostly moderate to mild and can sometimes be completely absent. Thus they are easily spread long distances through vegetative planting materials, although efficient seed transmission is also known for some species. Horizontal transmission has been reported by various mealybug or aphid species depending on the virus species. Some pararetroviruses, including some badnaviruses, can be present as integrated sequences in the genomes of some host plants termed endogenous para-retroviruses (EPRVs). Whereas such sequences are often fragmented and unable to reconstitute an infective viral genome, some EPRVs can be reactivated by certain stress conditions and form actively replicating viruses, a situation that occurs e.g. with certain Banana streak viruses in some bananas (16). Integration takes place through illegitimate recombination and is not necessarily associated with infection by a replicating virus. Southern blot analysis, immune-capture RT-PCR and rolling circle amplification are some of the techniques that have been employed to distinguish EPRVs from episomal viruses.

The aim of our study was to investigate in more detail some aspects of SPPV infecting sweetpotatoes, including its complete genome structure, how common it is, if SPPV like sequences can be found integrated into the host genome, if it can be transmitted to other plants, and if it is synergized by the SPCSV and or SPFMV.

## Results

### SPPV viruses are highly variable and ubiquitous among sweetpotato accessions

Entire genome sequences of sweetpotato pakakuy virus variants A and B were completed and found to be 7380 and 7961 nt in length respectively. There genomic structure was very similar to that typical of Badnaviruses, except that ORF3 was separated into two halves, which we designated ORF3a and ORF3b. They contain the movement (MP) and coat protein (CP) domains, or the asparyl protease, reverse transcriptase (RT) and RNaseH (RH) domains respectively and are separated by a short non-coding region. This region is highly variable between the viruses and was sequenced several times from independently amplified and cloned PCR products to ensure accuracy. For both viruses ORF3b is extended prior to the first methionine codon to overlap partially with ORF3a (12 and 21 nt respectively in SPPV-A and SPPV-B) and is found in a +1 reading frame as compared to ORF3a. SPPV-A and B share 79.5% nt identity over the complete genome and shared the same tRNA-met like region (TGG TAT CAG AGC GAG TAT) followed by a short stem-loop (GGC AGG CTA AGC CTA CC) and a putative leader sequence with extensive secondary structure (Fig 1).

**Fig 1.**
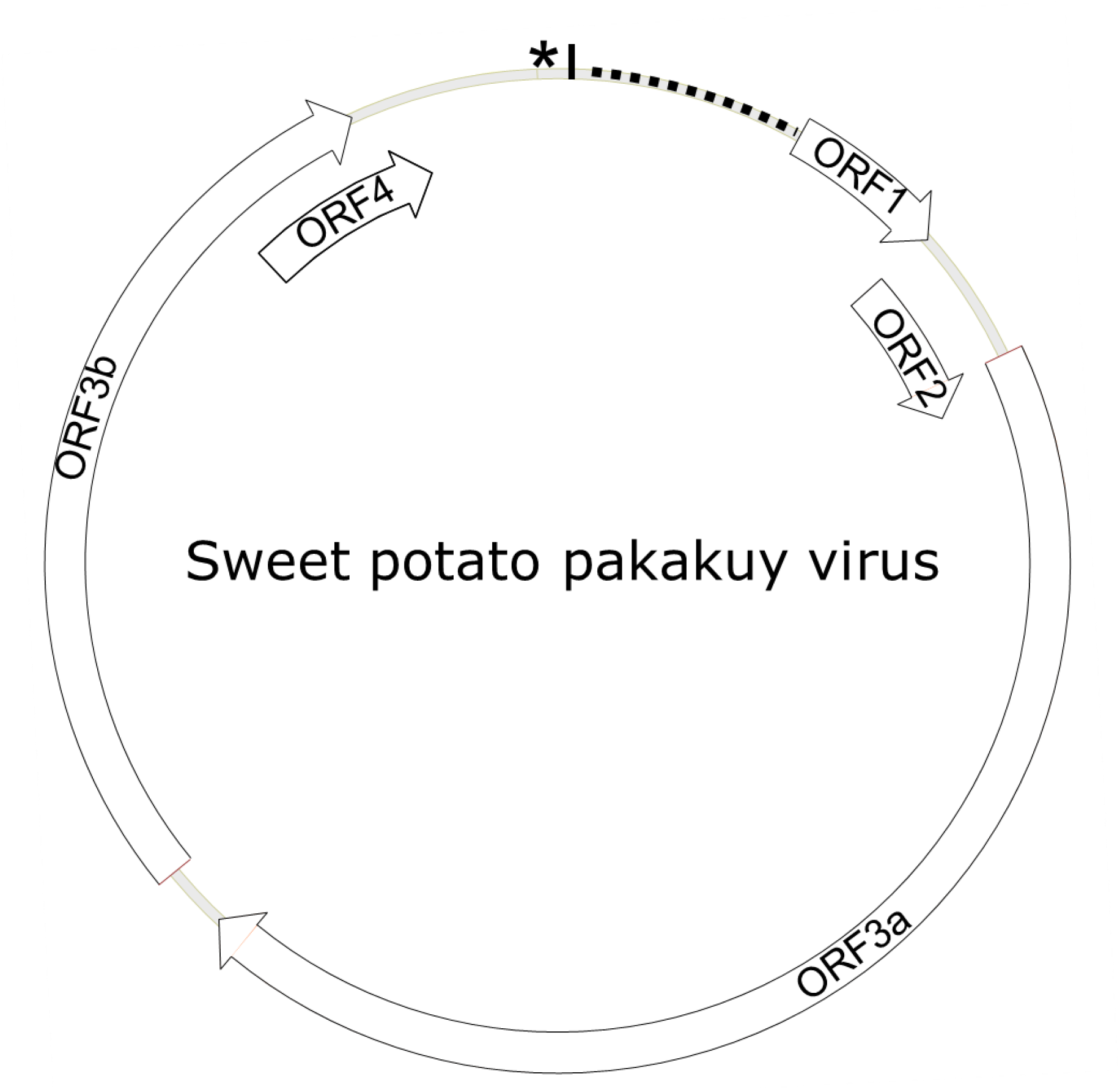
Genome structure of SPPV. Diagram depicting the genome structure of Sweet potato pakakuy virus (SPPV). Circle indicates the genome with box arrows indicating the locations of predicted open reading frames (ORFs) and numbered in order of occurrence. Star and vertical black line indicate the location of the tRNA-met like region and short stem-loop structure respectively, while the dotted line indicates location of a predicted leader sequence.

To determine how common these badnaviruses were among sweetpotato germplasm, we screened a collection of 78 sweetpotato genotypes from diverse geographic regions available in CIP’s germplasm collection with primers specific to SPPV-A and -B (Table 1) and found that many genotypes were infected by at least one of these viruses (Table 2). Subsequent siRNA deep sequencing and assembly of bulked RNA extracts which included samples recently received from Africa, produced additional contigs corresponding to badnaviruses, some of which were clearly distinct from SPPV-A and SPPV-B (S1 Data). Based on alignments of the RT and RH domains of the various sequences obtained, degenerate primers were designed and used to amplify the corresponding region from a subset of the 78 sweetpotato accession but also including 5 samples from African germplasm (Table 2). Phylogenetic analysis of alignments of nt or aa sequences of the RT or RT-RH domains resulted in a phylogenetic tree with three distinct and strongly supported clades, irrespective of the evolutionary inference method used, and a third more variable group, with less consistent support between phylogenetic inference method and/or nt substitution model applied (not shown). Two of the clades corresponded to SPPV-A and B, whereas the new clades were designated C and D (Fig 2). Whereas clades A-C were rather homogenous with mean within group nt variation of 1.1-2.2%, clade D was more variable with a mean variability of 10.5% and identifiable sub-groupings. Inclusion of additional sequences corresponding to SPPV from the GenBank did not affect the grouping into these clades (data not shown).

**Fig 2.**
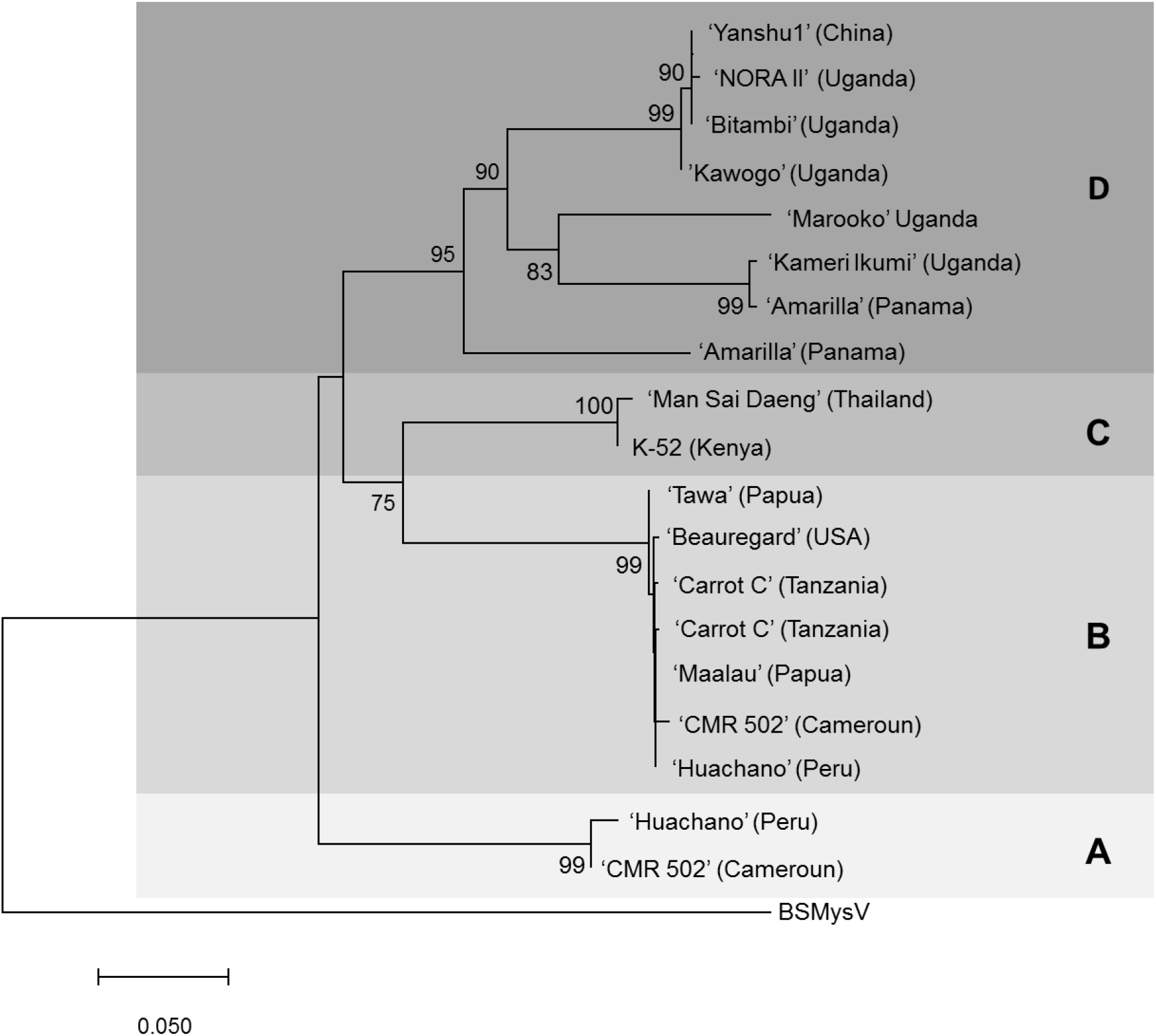
Phylogenetic tree of SPPV sequences covering the Reverse transcriptase and RnaseH domains amplified from sweetpotato accessions from around the world. The evolutionary history was inferred by using the Minimum Evolution methodand the evolutionary distances were computed using the Maximum Composite Likelihood method and are in the units of the number of base substitutions per site.. The optimal tree with the sum of branch length = 1.15686856 is shown. and is drawn to scale, with branch lengths in the same units as those of the evolutionary distances used to infer the phylogenetic tree. The percentage of trees in which the associated taxa clustered together is shown next to the branches based on 500 bootstrap replications when larger than 70%. The ME tree was searched using the Close-Neighbor-Interchange (CNI) algorithm at a search level of 1. The Neighbor-joining algorithm was used to generate the initial tree. The analysis involved 2o nucleotide sequences. All ambiguous positions were removed for each sequence pair. There were a total of 828 nt positions in the final dataset. Evolutionary analyses were conducted in MEGA7 (28). Isolates are indicated by the name of the variety from which they were amplified and the origin of the variety is provided in brackets for each of them. BSMysV (Banana streak mysore virus) was used as an outgroup for phylogenetic tree construction. Four phylogenetic groupings A, B, C and D are highlighted in blue, red, green and yellow respectively.

**Table 1.**
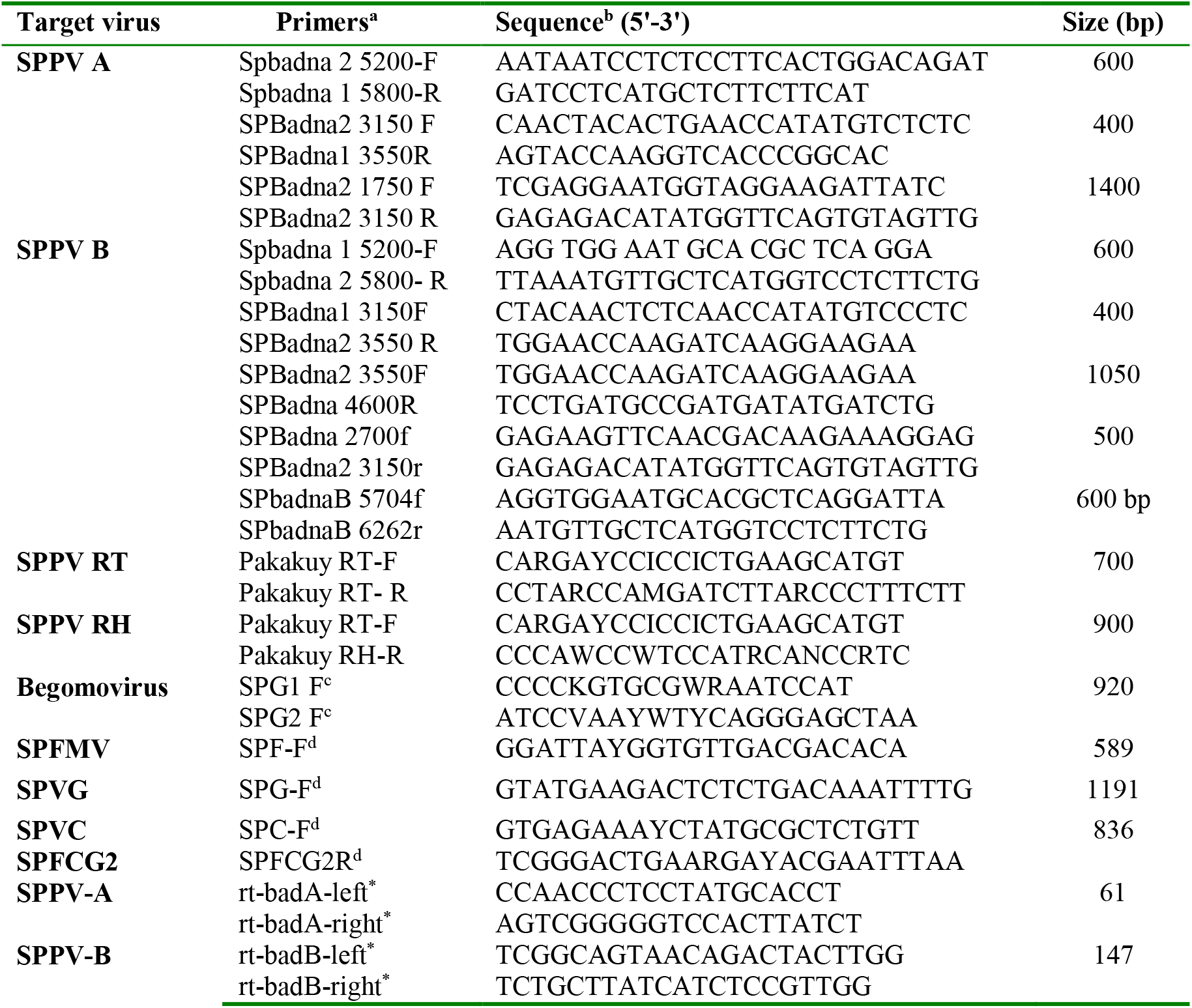
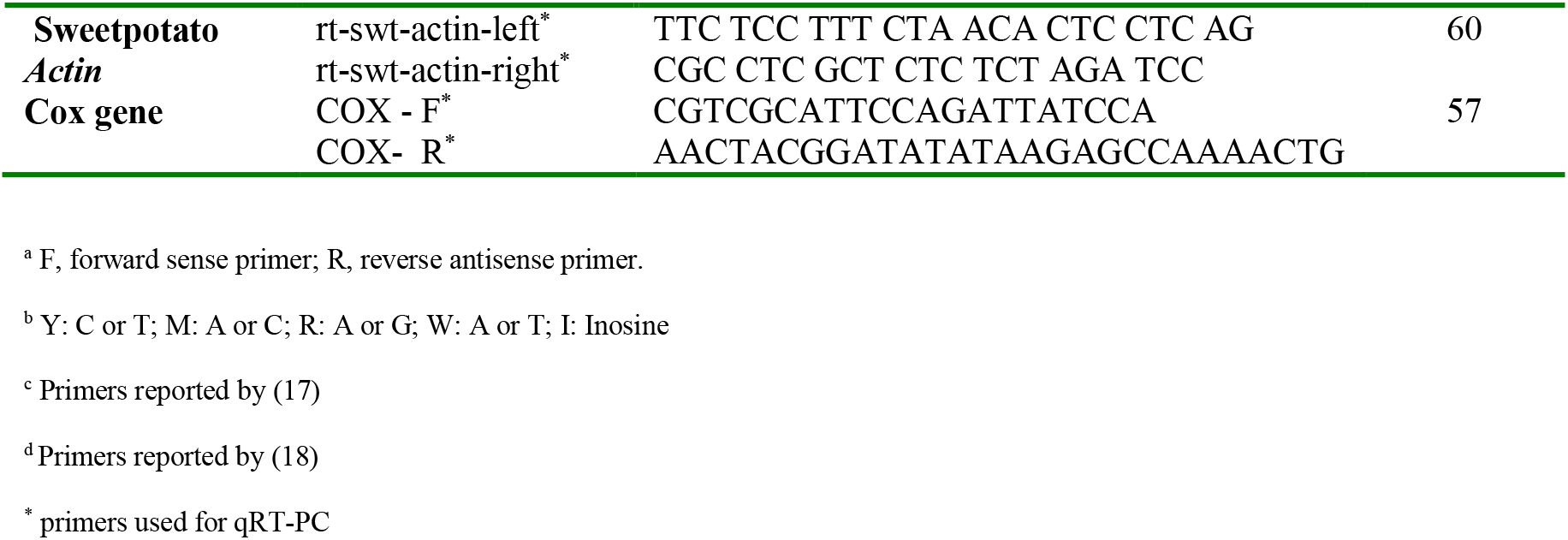
List of primers used in the detection for SPPV and qRT-PCR

**Table 2.**
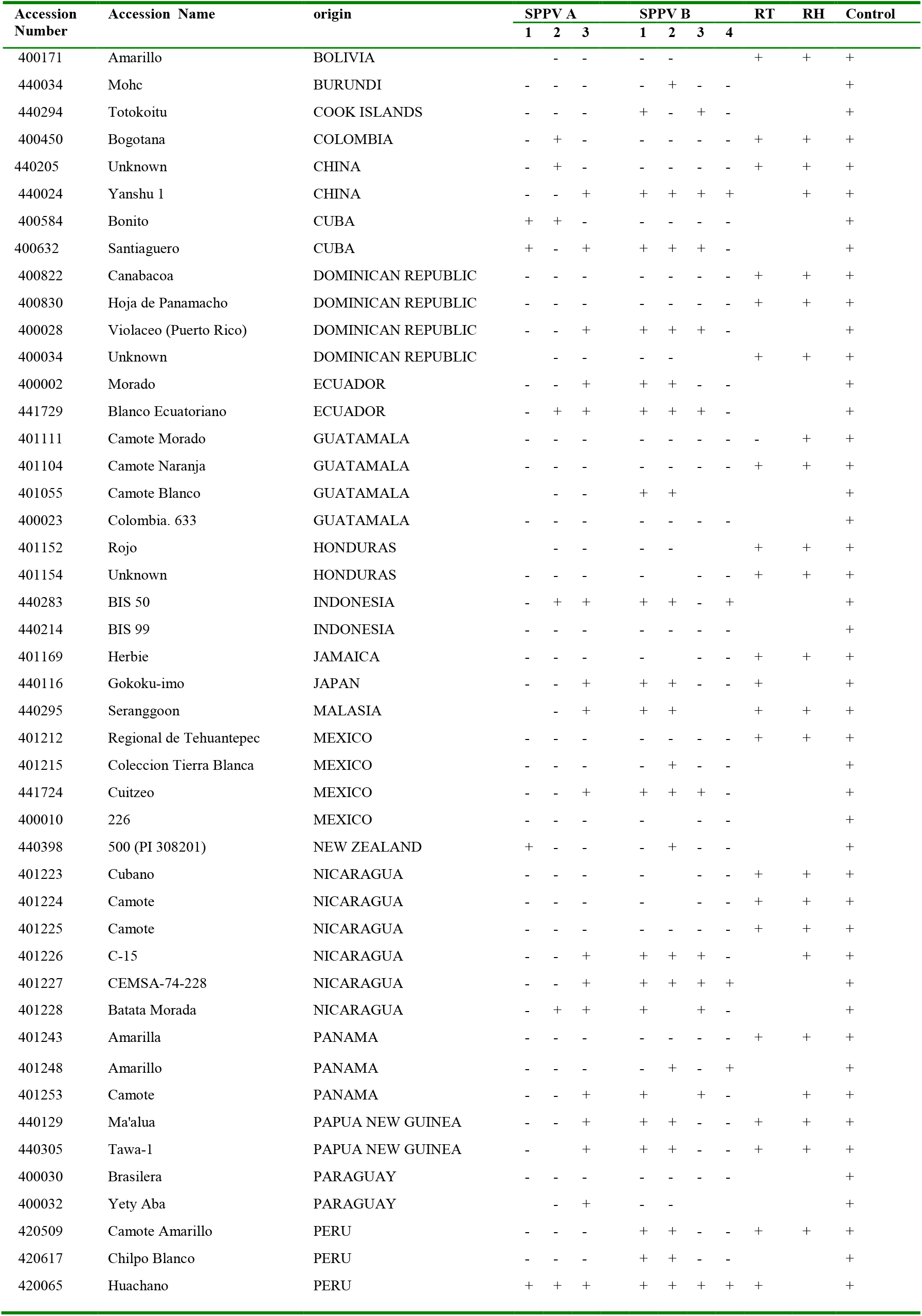
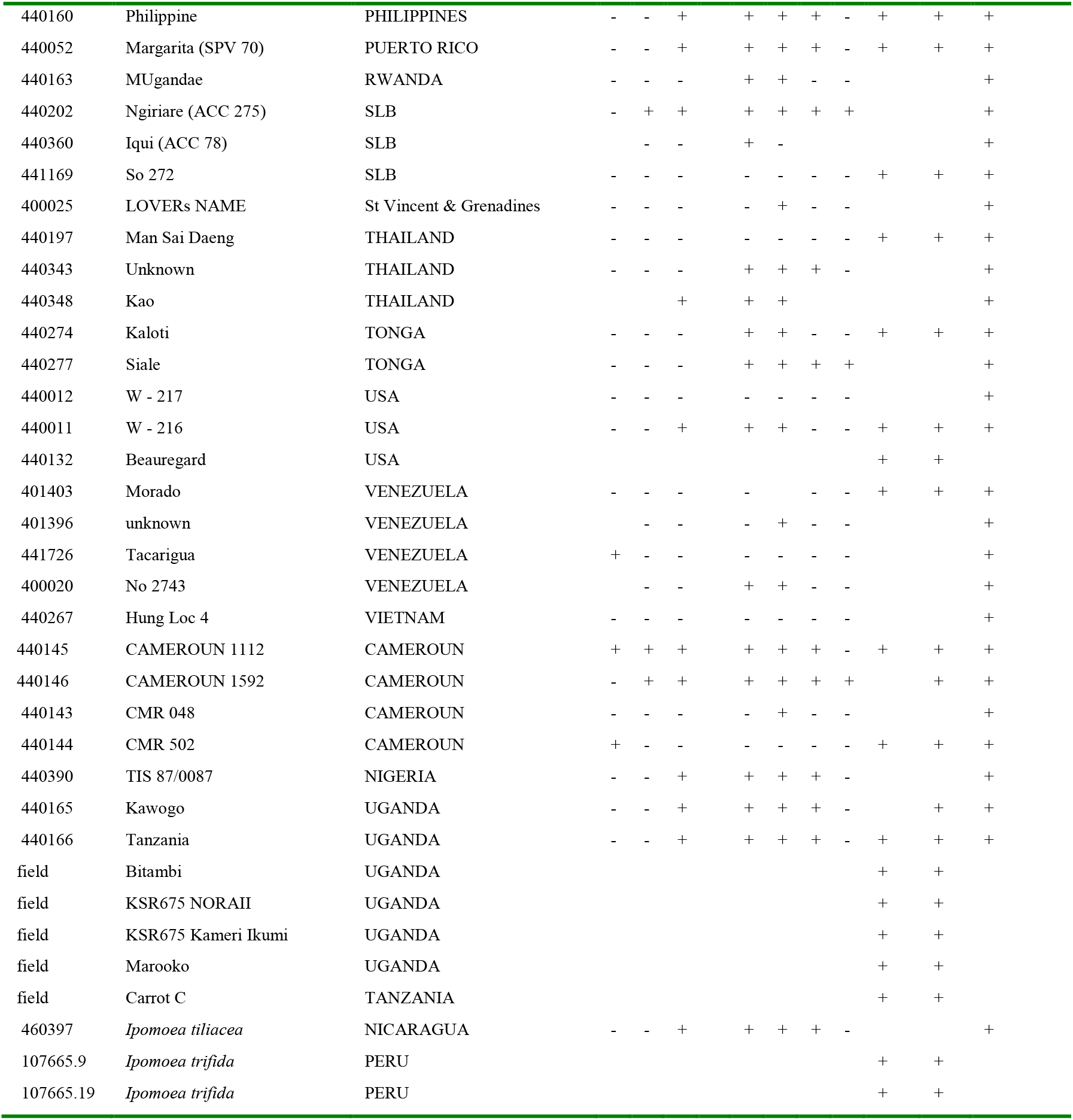
List of accessions used in this study and results of PCR tests for specific regions of SPPV-A and -B and degenerate primers.

### SPPV can be graft transmitted to indicator plants

Grafting experiments from sweetpotato (cv ‘Huachano’ infected with SPPV-A and -B) to sweetpotato (cvs ‘Man Sai Deng’ infected with SPPV-C and ‘Amarilla’ infected with SPPV-D, but which were not infected by SPPV-A or B) and from sweetpotato to *I. setosa* followed by PCR of the grafted plants resulted in positive reactions in some cases (treatment 4, 5, 9 and 13 in Table 3) indicating that SPPVs could be transmitted through this means although, apparently not with 100% efficiency, since in most cases only SPPV-B was transmitted (treatment 4, 5 and 9 in Table 3), whereas neither virus was transmitted to either sweetpotato cultivar when the source plant ‘Huachano’ was also infected by SPFMV and SPCSV (treatment 10 & 13, Table 3). To ensure that the virus detected in the graft inoculated *I.setosa* did not represent passively carried particles, the PCR positive *I.setosa* plants were used to graft inoculate a second *I.setosa*, which subsequently became PCR positive upon testing, except when the *I.setosa* was also infected by SPFMV and SPCSV (treatment 6 & 7 respectively in Table 3). In none of the cases were any visible symptoms discerned, except those of SPVD when the combination of SPFMV + SPCSV was included in the treatment, which are extremely severe in *I.setosa*. Cloning and sequencing of the PCR fragments from the serially inoculated *I.setosa* plants, confirmed they were identical to the sequence in the originally grafted plant in all cases.

**Table 3.** Results of graft transmission experiments

| Treatment | plants | PCR results* |  |  |
| --- | --- | --- | --- | --- |
|  |  | SPPV-A | SPPV-B | RT |
| 1 | Huachano<sup>1</sup> | 1/1 | 1/1 | 1/1 |
| 2 | Huachano (SPVD)<sup>1</sup> | 1/1 | 1/1 | 1/1 |
| 3 | I.setosa<sup>2</sup> | 0/2 | 0/2 | 0/2 |
| 4 | I.setosa + 1<sup>3</sup> | 0/2 | **2/2** | **2/2** |
| 5 | I.setosa + 2 | 0/2 | **2/2** | **2/2** |
| 6 | I.setosa + 4 | ND | ND | **2/2** |
| 7 | I.setosa + 5 | ND | ND | 0/2 |
| 8 | Man Sai Deng<sup>2</sup> | 0/2 | 0/2 | **2/2** |
| 9 | Man Sai Deng + 1 | 0/2 | **2/2** | 2/2 |
| 10 | Man Sai Deng + 2 | 0/2 | 0/2 | 2/2 |
| 11 | Amarilla<sup>2</sup> | 0/2 | 0/2 | **2/2** |
| 12 | Amarilla + 1 | **2/2** | **2/2** | 2/2 |
| 13 | Amarilla + 2 | 0/2 | 0/2 | 2/2 |

\*number of PCR positive plants/number of plants tested.
<sup>1</sup>source plants.
<sup>2</sup>test plants before grafting.
<sup>3</sup>test plant 25 days after graft inoculation with indicated source plant.

### SPPVs are seed transmitted in sweetpotato

A previously generated in-vitro germinated population from a cross between the cultivars Beauregard and Tanzania (19) which were both infected by SPPV (Table 2 & 4) were tested by PCR for presence SPPV in the established in-vitro plants and 76 out of 76 tested plants were found to be positive. PCR fragments were sequenced from ‘Beauregard’ (the mother), as well as three progenies and those of the progeny were found to be >99% identical to those found in Beauregard, and which corresponded to SPPV-B. In contrast all seedlings (203 plants) tested negative by PCR for begomoviruses, which both parents were also infected with, and also were PCR negative for SPFMV, sweet potato virus G (SPVG) and sweet potato virus C (SPVC), which were infecting the parent ‘Beauregard’ (Table 4). Thus, SPPV was transmitted to seed at very high efficiency.

**Table 4.** Results of PCR testing of in-vitro germinated seedlings and their parents

| *Virus identified* | *SPPV* | *Begomovirus* | *SPVG* | *SPVC* | *SPFMV* |
| --- | --- | --- | --- | --- | --- |
| Beauregard | 1/1* | 1/1 | 1/1 | 1/1 | 1/1 |
| Tanzania | 1/1 | 1/1 | 0/1 | 0/1 | 0/1 |
| B x T seedlings | 76/76 | 0/203 | 0/203 | 0/203 | 0/203 |

\*number of PCR positive plants/number of plants tested.

### Viral titers of SPPV are less than one copy per cell

Southern or dot-blot experiments using SPPV-A or -B specific chemi-luminiscent or radioactive probes consistently failed to detect either virus in several sweetpotato accessions tested irrespective if the plant was healthy, or infected by SPCSV, SPFMV or both viruses (data not shown). On the other hand sweetpotato DNA spiked with plasmid DNA containing the SPPV-A or -B probe fragments at a concentration corresponding to one or half a copy per sweetpotato genome were readily detected in Southern blot (Fig 3), indicating that the titers of these viruses must be well below these concentrations, and simultaneously imply these viruses are not integrated into the genome.

**Fig 3.**
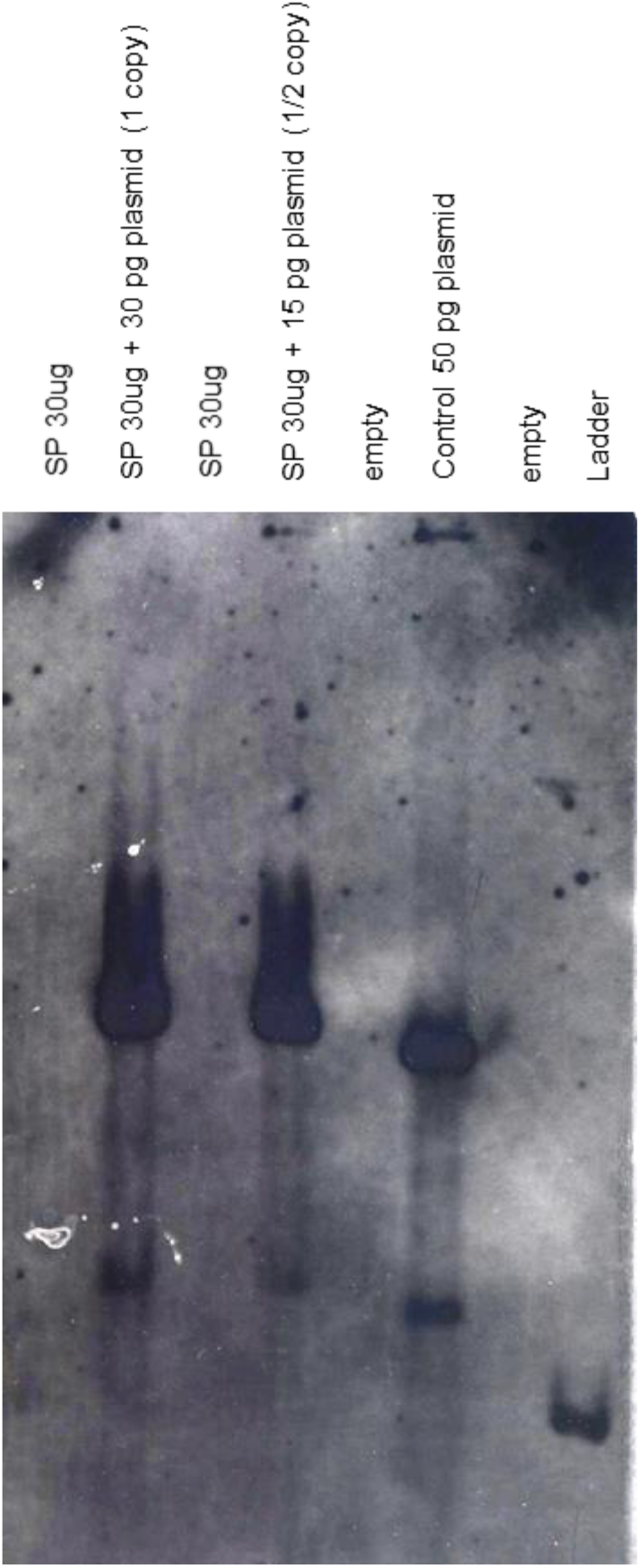
Southern blot of ‘Huachano’ DNA linearized with *PstI* and hybridized with a probe corresponding to SPPV-B. From left to right, the first and third lanes contain 30 ug of sweetpotato (SP) DNA, and the second and fourth lanes contain 30 ug of SP DNA spikes with 30 and 15 pg of plasmid (containing SPPV-B DNA fragment corresponding to the probe) respectivily corresponding to 1 or ½ a copy per sweetpotato genome equivalent; the 5^th^ and 7^th^ lanes are empty whereas the 6^th^ lane contatins 50 pg of SPPV-B plasmid DNA and the last lane a DNA ladder.

### SPPV titers are extremely low, are only minimally affected by co-infection of SPCSV and SPFMV whereas corresponding siRNA change their size distribution and are more abundant in SPVD affected plants

Because SPPV was below the detection limit of the Southern blot or dot-blot methods, a quantitative real-time PCR assay was developed to evaluate the distribution of virus titres in different leaves of sweetpotato cv Huachano. Results revealed qRT-PCR C(t) values averaging around 6 cycles below those of the reference gene *actin*, indicating extremely low concentrations in the extracted leaves (i.e. ~1% compared to actin). SPPV RNA concentrations between different leaves on the same plant showed up to 6 fold differences with the upper leaves tending to have higher titres (Fig 4). On the other hand, when comparing relative expression levels of virus infected plants to those of healthy plants, a significant increase of around 2.5 fold could be identified only for SPPV-B in plants infected with both SPCSV and SPFMV (Fig 4). Mapping of siRNA sequences determined from the three plants indicated that this correlated with increased siRNA production corresponding to SPPV-B viruses in plants infected by SPFMV and SPCSV as compared to other plants, mainly of 22 nt size, whereas 24 nt siRNAs were strongly reduced. Whereas this effect could be appreciated also in SPPV-A, it was slightly less extensively targeted by siRNAs than SPPV-B (Fig 5).

**Fig 4.**
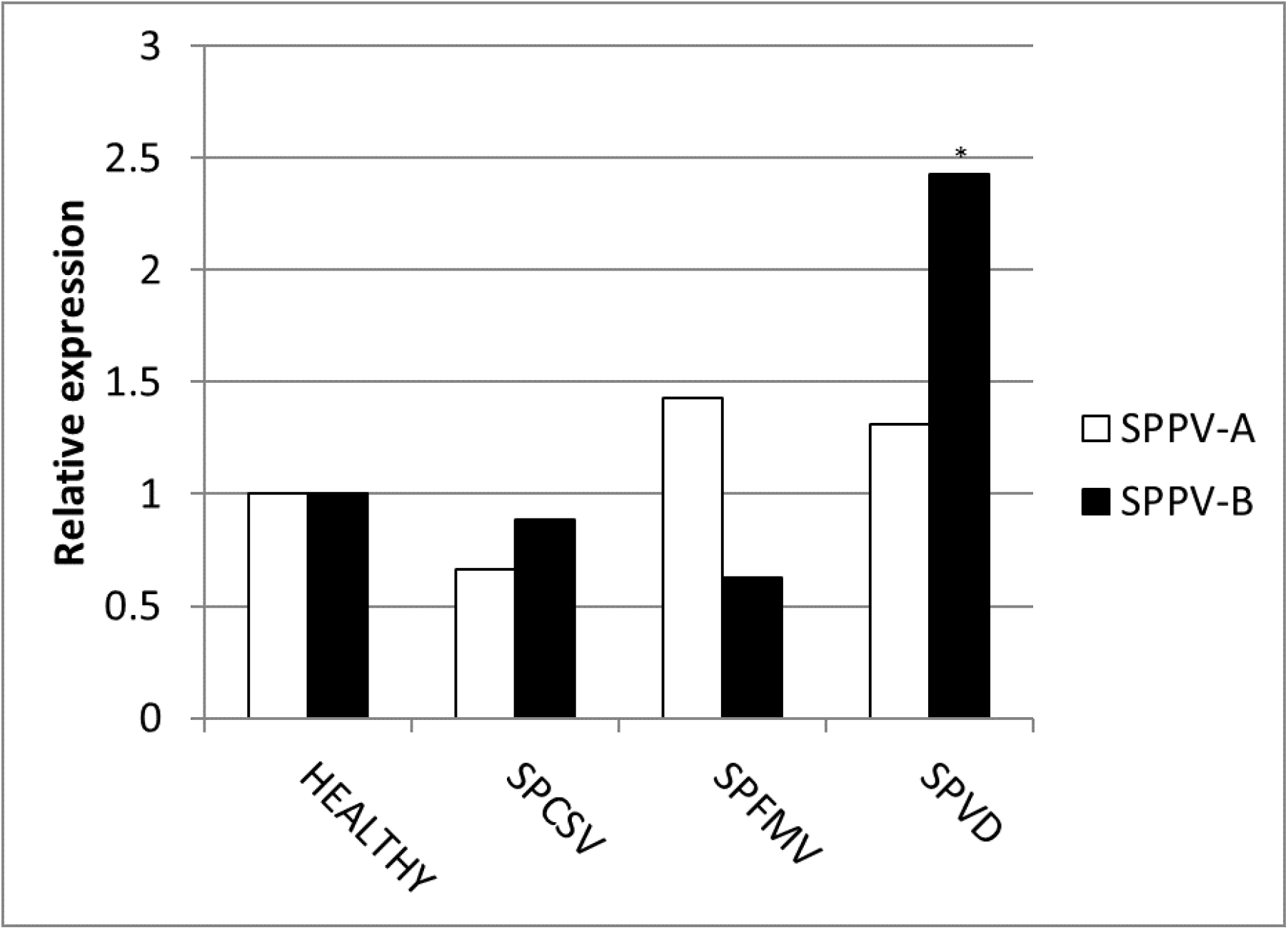
Relative expression for Badnavirus A and B. Graphic depicting the expression of SPPV-A and SPPV-B in leaves in co-infection with SPFMV, SPCSV or both viruses (SPVD) relative to a singly infected plants (healthy). * Significantly upregulated as compared to singly infected plants (healthy; p=0.001)

**Fig 5.**
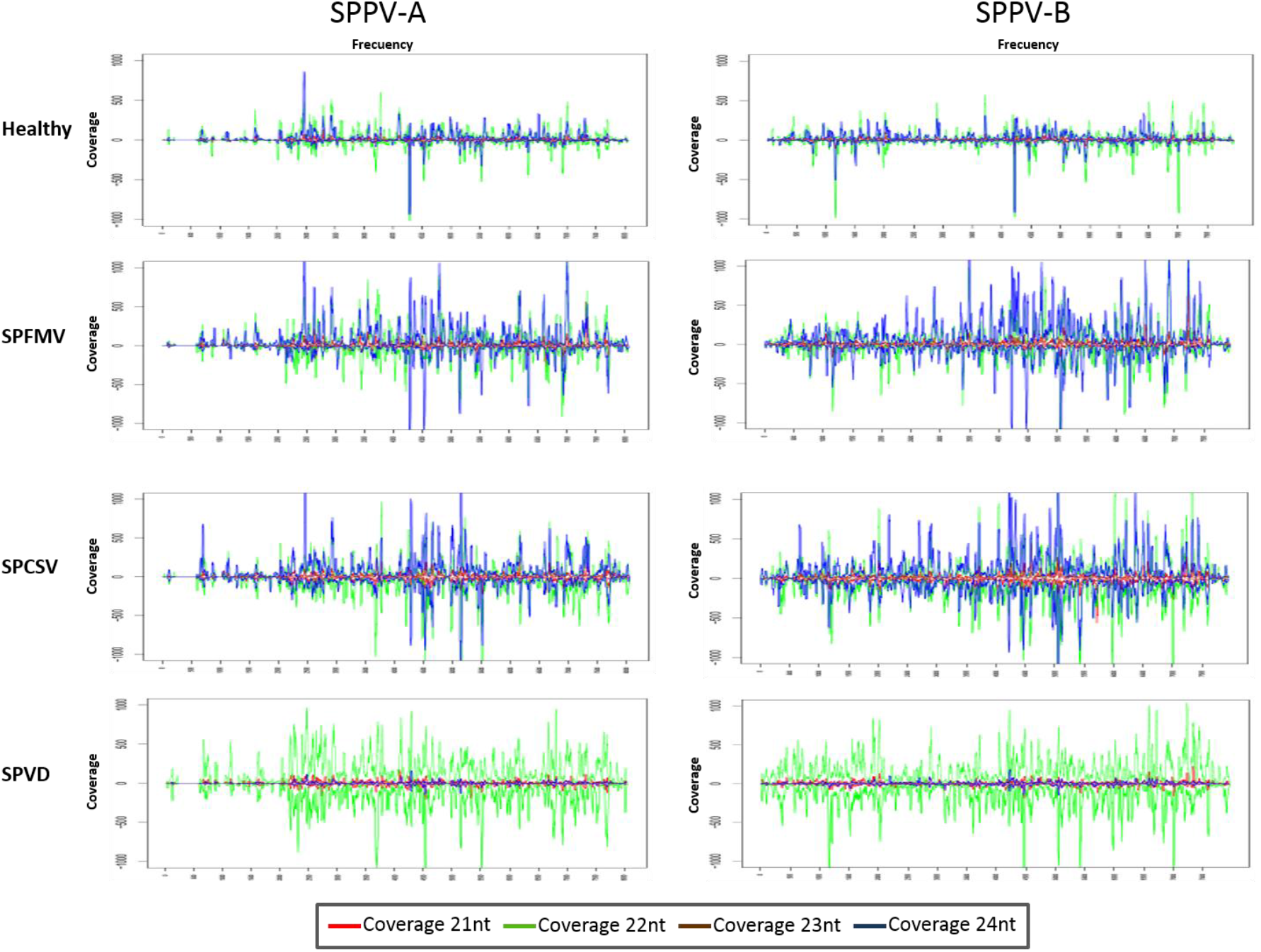
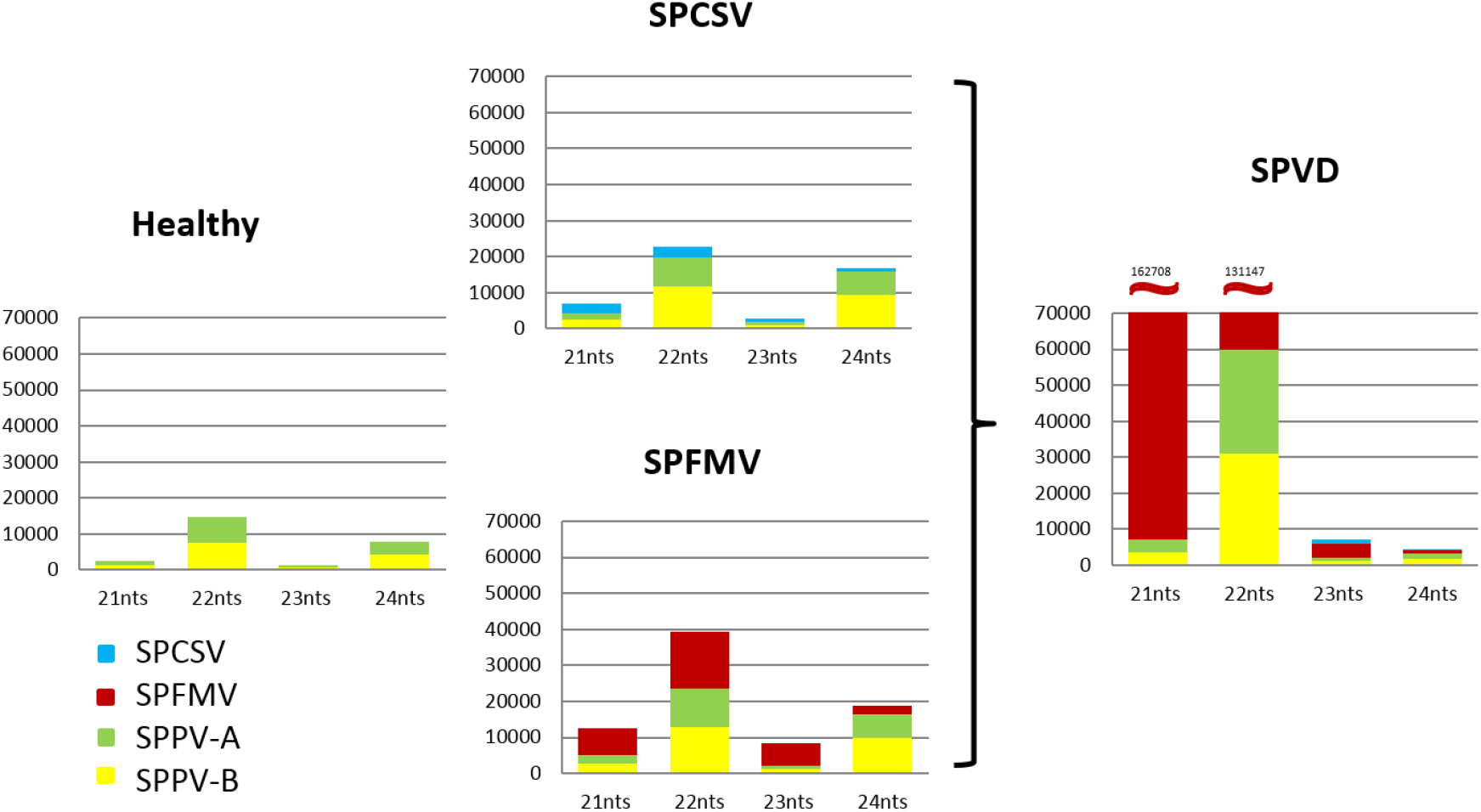
Size and distribution and quantities of siRNAs targeting SPPV in sweetpotato plants co infected with different viruses. A) Graphics show the normalized distribution (per million siRNA reads sequenced) of siRNA covering the genomes of SPPV-A (left) and -B (right) in healthy, SPFMV, SPCSV or dually (SPVD) co-infected plants. Horizontal axis indicates the nucleotide position of the virus wheras the verticle axis indicates the coverage of each nt position by siRNA sequences in sense (positive values) and antisense (negative values) orientation. Lines in red, green, brown and blue represent 21, 22, 23 and 24 nt siRNAs respectivily. B) Bar graphics showing the normalized (per million siRNA reads sequenced) quantity (vertical axis) and size (horizontal axis) of virus specific siRNAs in plants co-infected with different viruses. Green, yellow, red and blue sections in the bars correspond to SPPV-A, SPPV-B, SPFMV and SPCSV respectively.

## Discussion

Badnaviruses in sweetpotato remain somewhat enigmatic. SPPV was initially identified through siRNA sequencing from apparently healthy plants thought to be virus free (9), and has since then been identified in several NGS (10, 12, 14) studies and by PCR using specific primers based on the initial report (11, 13). Indeed, in this study we found that every plant we tested eventually turned out PCR positive for SPPV when degenerate primers were employed. However, results were not always consistent over time in all plants, a plant could test positive for a leaf sample at one time and negative at others (data not shown), suggesting low and unequally distributed concentrations in the plant. Nevertheless, because some badnaviruses are known to exist as EPRVs and EPRVs are also targeted by siRNAs through RNA silencing (16), it was important to confirm that what we were detecting were not integrated sequences. Our Southern blot experiments in Huachano unequivocally show that SPPV is not integrated in the genome of at least that cultivar and that SPPV concentrations are so low that they cannot even be detected by chemiluminescent hybridization. This conclusion was supported by qRT-PCR results showing that expression of SPPV RNA was around a hundred fold lower than that of the Actin reference gene (and a ~500 fold lower than COX reference). Sequence analysis of some of the amplified fragments from plants originating from different parts of the world showed considerable sequence variation between SPPV found in different genotypes, but also that many genotypes were infected by more than one variant, just like we found in cv. Huachano. This result suggests SPPV is an actively evolving virus.

Our virus transmission experiments also clearly showed SPPV-A and B could be transmitted by grafting to *I. setosa* and other sweetpotato plants infected with SPPV-C or D. It is noteworthy that in most cases only SPPV-B was transmitted, whereas qRT-PCR results suggested titres of both viruses were very similar. Perhaps SPPV-B is more adept at establishing infections than SPPV-A in a competitive situation. However the fact that SPPV-A and -B were found together and that SPPV-B could be transmitted to plants infected with SPPV-C or -D provided further evidence these viruses are not mutually exclusive. On the other hand co-infection of the source plant with SPFMV and SPCSV eliminated graft transmission of either virus to other sweetpotato plants, and serial transmission to *I. setosa*. SPVD is a severe disease in sweetpotatoes and sometimes lethal in *I. setosa*. It is conceivable that the stress caused by SPVD affects the formation of graft unions and other physiological factors that may impede efficient transmission of a virus already in such low titres.

Qin (2016) reported graft transmission of SPPV-A to *I. setosa* as determined by PCR, resulting in mosaic symptoms. However, it was not clear from that report if other viruses were infecting the original sweetpotato plants, and mosaic is not a typical symptom produced by badnaviruses. Indeed, this contrast with our findings which could identify no symptoms in *I. setosa* after graft transmission. None of the plants tested in this study showed any clear virus symptoms (except when affected by SPVD); the extremely low virus titres determined by qRT-PCR in the accession Huachano, suggest only very few cells might be infected and virus expression could be too low to induce any significant physiological changes in the plant that might manifest themselves in symptoms. However, without the availability of a plant lacking SPPV sequences it will remain impossible to determine any biological impact SPPV may have on sweetpotato production. Our qRT-PCR and siRNA sequencing experiment in plants co-infected with SPFMV and SPCSV indicated only minor effects on SPPV titres, which were only significant in the case SPBaV-B in dual infection with SPFMV and SPCSV. The modest 2 fold increase observed, however, seems unlikely to be able to mediate much impact, particularly when considering the several hundreds of fold increase of SPFMV caused by SPCSV co-infection (5, 7, 20). In contrast, as had been previously observed (9), infection of both SPFMV and SPCSV had a marked effect on the amount and size of siRNAs targeting SPPV, but also infection by SPFMV and SPCSV alone affected siRNA amounts (but not size). These changes can probably be attributed to the effects of expression of the different silencing suppressors of both viruses, but as evidenced from qRT-PCR experiments, these nevertheless had minimal effect on SPPV titres themselves.

The genomes organizations of the two SPPV isolates determined in this study are slightly different from other badnaviruses in that ORF3 is divided into two (3a and 3b), a situation also found in cassava vein mosaic virus (genus *Cavemovirus*). Although ORF3b may be expressed from a separate mRNA the possibility remains that it is expressed through +1 ribosomal frameshifting as there is an overlap between the two ORFs when extending ORF3b 5’of it’s first potential initiation codon.

Because ‘Huachano’ plants originated from in-vitro plants that had been submitted to thermotherapy and meristem tip culture for virus elimination, it suggests that despite its low virus titers SPPV is able to maintain itself in meristematic tissues. Indeed, attempts in other laboratories to eliminate viruses by thermotherapy and meristem excision failed to eliminate SPPV (Christopher Clark, personal communication). On the other hand several accessions of a wild sweetpotato relatives, *I. tiliacea* and *I.setosa*, which are grown from seed, were also found to be positive suggesting that the virus could also be transmitted by seed. Seed transmission was confirmed to be highly efficient in sweetpotato by testing in-vitro germinated seedlings derived from a cross between ‘Beauregard’ and ‘Tanzania’, whereas other viruses infecting either parent showed no evidence of seed transmission, as expected. Perhaps this is the principal mechanism by which SPPV has maintained and spread itself among sweetpotatoes worldwide as it seems hard to imagine any vector could be very efficient at transmitting SPPV between sweetpotatoes when titres are so low. On the other hand, the sequence variation found between different genotypes indicates they are not all descending from the same source and it could be possible that sweetpotato is occasionally (re-)infected from an unknown source plant. Electron microscopic studies by Sim et al.,(21) claimed to identify badnavirus like particles in *Ipomoea nil* plants and it could be interesting to survey more wild *Ipomoeas* spp. as possible sources of SPPV.

Based on their apparent universal presence in sweetpotatoes and lack of obvious symptoms and vertical transmission over generations, SPPV could be considered among the persistent (or criptic) viruses (22, 23). Previously identified persistent viruses have been exclusively RNA viruses belonging to specific families like Partitiviridae & Totiviridae (dsRNA) or Endornaviridae (ss+RNA). Persistent viruses are characterized by vertical transmission, from seed and or pollen and cell-to-cell by redistribution in dividing cells; they lack movement proteins and in the case of endornaviruses even lack any discernible proteins besides the replicase. Because they also lack any discernible symptoms in infected plants they have been considered commensal or mutualistic in their interaction with plants, although mutualistic interaction have only been proven in a couple of cases (24, 25). Whether the presence of SPPV in all the genotypes we tested may similarly results from a mutualistic interaction or even a process of human selection remains to be determined but is certainly an intriguing possibility.

## Materials and Methods

### Plant material and viruses

Plant materials used are summarized in Table 1. A total 78 accessions from the worldwide sweetpotato collection (including five newly acquired accessions not yet assigned accession numbers) and three related wild *Ipomoea* spp. at the International potato center (CIP) genebank were evaluated by PCR for presence of SPPV. They were established and maintained in an insect-proof greenhouse at 27±1°C at CIP as a backup to the in-vitro collection since their original acquisition. cv ‘Huachano’ used in this study originated from in-vitro ‘virus free’ plants that had passed through thermotherapy and meristem tip culture (9). A mapping population of a cross between cv Beauregard and Tanzania was described previously (19). Plants of the universal sweetpotato virus indicator *I. setosa* and one accession of *I. tiliacea* were grown from seed produced at CIP virology unit.

### Nucleic acid extractions

Total DNA from infected *Ipomoea* spp. leaves was extracted using the CTAB method (26). Leaf tissue (approximately 250- 400mg) was ground to a fine power in liquid nitrogen using a mortar and pestle, in the presence of 2ml of extraction buffer, followed by an incubation period at 60°C for 30 min and addition of an equal volume of chloroform: isoamyl alcohol (24:1). The homogenate was vigorously shaken at room temperature for 10 min using a vortex and after centrifugation at 12000 g for 10 min, the supernatant (~500ul) was recovered, mixed with same volume of Isopropanol and centrifugated at 12000 g for 10 min. The precipitated DNA was washed with 70% ethanol, dried, resuspended in 100 ul of Nucleases free water (NFW), and kept at – 20°C until analysis.

Total RNA was extracted using CTAB RNA method modified with LiCl (Adapted from (27)), from fresh leaves by grinding tissue with a hand roller, adding 10x (v/w) of CTAB buffer followed by centrifugation in a microfuge at maximum speed for 5 min at room temperature. Subsequently an equal volume chloroform IAA (24:1) was added and the homogenate was mixed thoroughly before centrifuging again at maximum speed in a microfuge for 5 min. The supernatant was carefully removed and mixed with an equal volume of 4M LiCl and left overnight on ice in fridge. The precipitated RNA was centrifuged for 20 min at maximum speed in a microfuge and washed with 70% ethanol, the pellet was dried and kept at – 70°C until analysis.

### PCR amplifications, sequencing and sequence analysis

PCR reactions were performed in a total volume of 25ul containing 2mM MgCl2, 1X PCR reaction buffer, 0.2mM dNTPs, 0.2 uM of each primer, 0.02units Taq DNA polymerase (Promega) and 1 ul (100ng) of DNA sample. DNA from healthy *I. setosa* plants was also included in these experiments as negative controls. PCR amplification of virus specific fragments of SPPV-A and –B from cv Huachano, was performed using primers designed based on previously reported partial sequences (9). Additional primers were designed based on the conserved functional domains present in the putative polyprotein encoded by open reading frame (ORF) 3 for detection SPPV-A and –B in germplasm and grafting experiments (Table 2). PCR was performed in a DNA thermal cycler (Applied Biosystems) with an initial denaturation cycle for 2 min at 94°C, followed by 35 cycles for 30s at 94°C, 30s at 56°C, 1 min at 72°C, and a final extension for 10 min at 72°C. The amplified products were loaded in a 1% agarose gel stained with GelRed™ (Biotum). Amplified fragment were cloned into pGEM-T Easy (Promega). Sequencing of PCR amplified fragments using the Sanger method was performed by Macrogen (Seoul, Korea)

Nucleic acid alignments and phylogenetic analysis were performed using Mega7 (28) (www.megasoftware.net) using maximum likelihood and the substitution models calculated to best fit the alignment data.

### Quantitative real-time PCR

Sweetpotato plants were infected with SPFMV, SPCSV, both viruses under controlled greenhouse conditions in Lima, Peru. Cuttings were taken from infected plants and grown for 3 months after which leaves were collected from basal, middle and top of each plant. Total RNA was extracted using CTAB as described above. 1 μg of total RNA was treated with 2 U of Turbo DNA-*free*™ (Ambion) in a total volume of 10 μl according to the manufacturer’s protocol. After heat deactivation of the DNase enzyme cDNA synthesis was carried out using 1ul of the DNase treated RNA, random primers (Invitrogen) and Superscript™ III reverse transcriptase (Invitrogen) in a total volume of 20 ul according to the manufacturer’s protocol.

The qPCR primers were for actin, SPPV-A and -B (Table 1) were designed using the “Primer3” open source bioinformatic tool (http://primer3.sourceforge.net/). Primers for cytochrome oxidase (Cox) have been previously reported (29).

The qPCR experiment was set up with three replicates per sample per plate. The Power SYBR^®^ Green PCR Master mix (Applied biosystems) was employed for the qPCR with 4 μl of cDNA solution in a volume of 10ul according to the manufacturer’s protocol. The reaction and the detection of the fluorescent signal were performed with the Mx 3005P qPCR System (Stratagene). Actin and Cox genes were used as internal control and reference genes for data normalization. The data analysis was carried out using the 2^(-ΔΔCt)^ method (30) to determine relative expression levels. The REST2009 software (Qiagen) was used to determine statistical significances in relative expression between different samples.

### Southern blots

A plasmid containing SPPV insert was used to synthesize non-radioactive probe using the PCR DIG Probe Synthesis Kit (Roche) with the primers SPbadnaB 5704f and SPbadnaB 6262r (Table 1) which amplified a ~600 bp fragment of ORF 3b region. The probe was amplified with a thermal cycler (Piko, Finnzymes) using 30 cycles, each consisting of 30 sec at 95°C, 30 sec at 60°C and 40 sec at 72°C. A final step of 7 min at 72°C also was included. Total DNA from sweet potato cv. ‘Huachano’ foliar tissue was extracted using CTAB method as described above. Extracted DNA (30ug) was digested with *Ecor* I and separated by 0.8% agarose gel electrophoresis in TAE containing GelRed™ overnight at 30v. The plasmid containing the SPPV insert linearized with *Pst* I and used as a positive control. After depurination, denaturation and neutralization steps, DNA was transferred to a positively charged nylon membrane and fixed with ultraviolet light treatment (UV Stratalinker 2400 Stratagene) DNA was then pre-hybridized, hybridized and developed with CDP-Star, ready to use kit (Roche) following the manufacturer’s procedures and Kodak Biomax light film (Sigma).

### Graft transmissions

*I. setosa*, and two sweetpotato genotypes (Amarilla and Man Sai Deng, CIP Germplasm accession numbers 401243 and 440197 respectively) which tested negative for SPPV-A or -B by PCR screening were selected for graft transmission experiments from cv Huachano. Plants were tested by PCR for SPPV-A and B and generic SPPV primers before graft inoculation, after which they were inoculated by side grafting a single node including leaf of the sweetpotato cv Huachano which was either healthy, or affected by SPVD. All plants were maintained in a greenhouse under controlled conditions at 27±1° C and monitored for symptoms up to 8 weeks and tested by PCR at 25 days post grafting. The success of the graft union was confirmed by survival of the grafted scion throughout the experiment. PCR fragments amplified by SPPV-A and –B specific primers were sequenced to corroborate the results. To confirm that positive PCR results in graft inoculated *I.setosa* plants were not due to passive transmission of virus from the grafted sweetpotato scion, serial transmission was performed by grafting scions from the first *I. setosa* plants to two new *I. setosa* plants. The serially grafted *I.setosa* plants were tested by PCR using the generic primers RT-F and RT-R at 21 days post inoculation.

### siRNA sequencing and assembly

To evaluate effect of co-infection of SPFMV and SPCSV on SPPV siRNA levels, leaves from the middle of 1 month old healthy, SPFMV, SPCSV or SPVD affected samples of ‘Huachano’ were used for RNA extraction. RNA was extracted using Trizol reagent according to the manufacturers instructions. RNA was then run in a 3.5% agarose gel and the band corresponding to siRNAs cut and purified using quantum prep gel purification columns (Bio-Rad). Purified siRNAs were sent to Fasteris Life Sciences (Switzerland) for sequencing on an Illumina Hiseq 2000. Small RNA sequences were downloaded and are accessible https://research.cip.cgiar.org/confluence/display/cpx/CIP.sweetpotato.2014, siRNA sequences were mapped against the genomes of SPPV-A and SPPV-B using MAQ and coverage of their respective genomes by siRNAs was visualized using a custom script (available from authors upon request).

To identify SPPV infecting sweetpotato cultivars collected from the field in Africa, RNA was extracted from leaves of seven different plants and combined with 13 additional samples from potato and other plant species from and processed and sequenced as described above (accessible from https://research.cip.cgiar.org/confluence/display/cpx/GAF13-14), except that sequences were de-novo assembled using velvet as described previously, and contigs were submitted to BlastX at NCBI selecting Badnaviruses as organism search set. The hit tables were downloaded and imported into Microsoft Excel for presentation (S1 Data), and contigs with hits were aligned to SPPV-A and -B sequenced for design of degenerate primers able to identify all SPPV variants.

### Accession numbers

The complete genome sequences of SPPV-A and SPPV-B determined from ‘Huachano’ were submitted to GenBank, receiving accession numbers FJ560945.1 and FJ560946.1 respectively. Partial sequences of the reverse transcriptase and/or RNaseH domains from addition cultivars received GenBank accession numbers KM000051-KM000054, KM009088-KM009100 & KM015301-KM015304.

## Acknowledgements

We thank Segundo Fuentes, Dora Quispe, Genoveva Rossel and David Tay for support and sharing of materials. We thank Jari Valkonen and Isabel Weinheimer for sharing primer sequences for real-time PCR.

---

**S1 Data. BLASTX results of contigs assembled from a mix of African sweetpotato cultivars with similarity to SPPV**. Contigs assembled from siRNA sequences of a bulk sample including several African sweetpotato cultivars with similarity to Badnaviruses (first sheet) and the hit table for each contig (sheet 2).

